# Controlling cell shape on hydrogels using lift-off patterning

**DOI:** 10.1101/111195

**Authors:** Jens Moeller, Aleksandra K. Denisin, Joo Yong Sim, Robin E. Wilson, Alexandre J.S. Ribeiro, Beth L. Pruitt

## Abstract

Polyacrylamide gels functionalized with extracellular matrix proteins are commonly used as cell culture platforms to evaluate the combined effects of extracellular matrix composition, cell geometry and substrate rigidity on cell physiology. For this purpose, protein transfer onto the surface of polyacrylamide hydrogels must result in geometrically well-resolved micropatterns with homogeneous protein distribution. Yet the outcomes of micropatterning methods have not been pairwise evaluated against these criteria. We report a high-fidelity photoresist lift-off patterning method to pattern ECM proteins on polyacrylamide hydrogels ranging from 5 to 25 kPa. We directly compare the protein transfer efficiency and pattern geometrical accuracy of this protocol to the widely used microcontact printing method. Lift-off patterning achieves higher protein transfer efficiency, increases pattern accuracy, increases pattern yield, and reduces variability of these factors within arrays of patterns as it bypasses the drying and transfer steps of microcontact printing. We demonstrate that lift-off patterned hydrogels successfully control cell size and shape and enable long-term imaging of actin intracellular structure and lamellipodia dynamics when we culture epithelial cells on these substrates.

## Introduction

Cell culture substrates patterned with extracellular matrix (ECM) are widely used to mimic the spatial organization and rigidity of the *in vivo* cell microenvironment *in vitro*. These cell culture platforms enable reductionist studies of the mechanobiology of healthy and diseased tissues under physiological-stiffness conditions ^1^. Specifically, polyacrylamide (PAAm) hydrogels are commonly used because these platforms can be functionalized with ECM and tuned in their mechanical properties to replicate different tissue stiffness ranging from ~0.1 kPa to ~40 kPa ^2^. Yet techniques for patterning proteins on PAAm have lacked quantitative assessment, which is critical for developing and comparing protocols to reliably restrict cells to user-defined shapes. The spatial resolution and accuracy of the protein patterns will directly impact the cellular response which is of particular importance for mechanobiological studies on the organization and force transduction within the actin cytoskeleton ^3^ and cellular adhesions ^4^.

Broadly, two main strategies exist to pattern ECM on PAAm gels (reviewed in ^5^): i) selective activation of the gels for covalent attachment of proteins to activated regions (e.g. direct surface functionalization using UV-reactive sulfo-SANPAH crosslinkers ^2^ or polymerizing Nhydroxyacrylamide into the hydrogel surface ^6^) and ii) co-polymerization of ECM proteins into the gels during gelation through direct contact of the acrylamide precursor mix with a protein-patterned coverslip. The first method, direct surface functionalization, uses expensive functionalization reagents and also depends on reagent quality and reaction time as the chemicals are unstable in aqueous media and in the presence of oxygen ^5^. The method of co-polymerizing ECM proteins relies on patterning glass coverslips with protein and placing them in direct contact with the hydrogel during polymerization. Although the detailed molecular mechanism of protein incorporation into the polymerizing gel is unknown, this method has successfully been applied to functionalize hydrogels with a variety of ECM proteins ^7-9^.

Protein patterns on glass coverslips are often created by microcontact printing (μCP) using elastomeric ‘stamps’ ^10^. μCP involves casting polydimethylsiloxane (PDMS) on microfabricated master structures created by photolithography to create stamps by replica molding ^11^. Most groups use μCP since PDMS casting and contact printing protocols are straightforward once the master structures on silicon wafers are made ^7, 12, 13^. However, μCP relies on the transfer of dried proteins from a deformable PDMS stamp and thus the accuracy, resolution, and pattern design are limited and critically depend on the PDMS stamp preparation and handling ^14, 15^. To improve the accuracy, alternative protein patterning methods have been developed. For example, dip-pen nanolithography enables direct writing of proteins on flat, solid substrates with nanometer precision ^16^. However, this method is serial and does not permit the functionalization of hydrogels. To overcome those challenges, protocols based on the selective oxidation of poly(l-lysine)-graft-poly(ethylene glycol) (PLL-g-PEG) layers by deep UV irradiation through a photomask or via projection lithography have been reported to pattern proteins on glass ^17, 18^. Such patterns can subsequently be transferred to a hydrogel ^9^ thereby decoupling pattern generation from hydrogel functionalization. Those methods however require either access to a collimated deep UV light source or a UV projection system, which are not readily available in most laboratories. Further, the PLL-g-PEG layer must either be dried prior to the UV irradiation, which requires a rehydration step prior to protein incubation ^17^, or a photoinitiator must be added during the UV exposure step that has to be removed completely to re-establish the biopassive properties of the adlayer ^18^.

In this work, we present a photoresist lift-off patterning (LOP) method to control the shape of cells on PAAm hydrogels with high fidelity. Our method integrates advances in: i) contact photolithography and photoresist lift-off widely used in the semiconductor and microfabrication industry ^19^, ii) the molecular assembly and patterning of biopassive PLL-g-PEG coatings on glass^20-22^, and iii) the protein transfer from glass to PAAm hydrogels ^12^.

We create protein-patterned glass coverslips by photoresist lift-off-assisted patterning of PLL-g-PEG and transfer the protein pattern to PAAm gel surfaces by co-polymerization. To demonstrate the utility of this approach, we successfully controlled the shape of MDCK cells cultured on patterned hydrogels and followed the cells’ cytoskeletal and membrane dynamics. We benchmark the LOP technique to the widely used μCP across a range of physiologically relevant hydrogel stiffness (5 kPa, 10 kPa and 25 kPa) and analyze the pattern accuracy and transfer efficiency from the glass to the PAAm gel. We find that the LOP protocol improves both the pattern transfer efficiency and the pattern accuracy, thereby reducing the pattern variability and increasing the predictability of the engineered *in vitro* cell culture models.

## Methods

### Photoresist lift-off assisted patterning of ECM proteins (LOP)

ECM patterned glass coverslips were fabricated by photoresist lift-off (see process flow in Figure 1, full protocol in Supplemental Material). We cleaned coverslips with acetone, isopropanol, and water, followed by thoroughly drying them on a hot plate. We then spin-coated S1818 photoresist (Microchem) on coverslips using standard contact photolithography and photopatterned the 2μm thick resist layer (40-50 mJ/cm^2^ at 365 nm, OAI Instruments) (Figure 1 i). Following plasma activation, we incubated the S1818 patterned substrates with 0.1 mg/ml (poly(l-lysine)-graft-poly(ethylene glycol) (PLL-g[3.5]-PEG (2kDa), SuSoS AG) for one hour. After photoresist lift-off in 1-methyl-2-pyrrolidone (NMP, Sigma 328634), we backfilled the PLL-g-PEG patterns with 100 μg/ml of Oregon Green-488 or Alexa Fluor 568-labeled gelatin solution in PBS pH 7.4 for 1 hour in the dark (Thermo Scientific, G13186, A10238) (Figure 1 ii-iv). The slides were washed thoroughly with DI water and excess liquid was removed by blotting on filter paper immediately prior to gel transfer. We chose gelatin, hydrolyzed collagen I, as a model ECM protein to mimic the epithelial basement membrane because the Arg-Gly-Asp (RGD) sequence critical for cell adhesion, migration and proliferation is preserved ^23^. Gelatin, in contrast to collagen I, is available commercially with fluorescent labels or can be functionalized with standard protein labeling kits.

**Figure 1.**
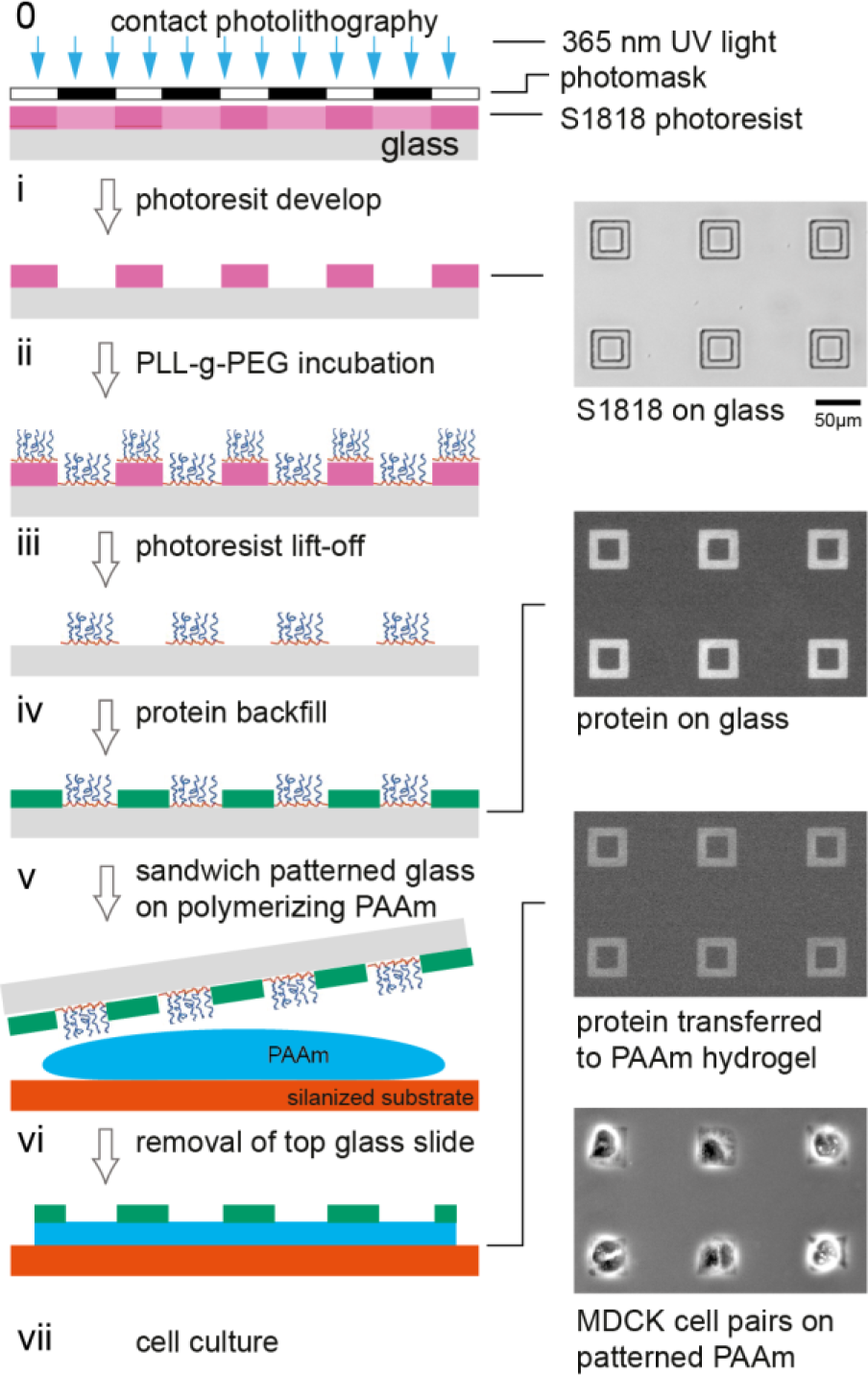
LOP fabrication of protein patterns on polyacrylamide gels. (i) Photoresist patterns are fabricated by standard contact photolithography on glass coverslips. Inset at right shows array of S1818 photoresist features after development. (ii) Unspecific protein adhesion to the resist-patterned coverslip is blocked by incubating with biopassive PLL-g[3.5]-PEG(2kDa) copolymer. (iii-iv) Following photoresist lift-off, the resulting PLL-g-PEG pattern is backfilled with the ECM protein of interest. Inset at right shows a fluorescence micrograph of labeled gelatin on glass after backfill. (v) To transfer the protein pattern to the PAAm gel, the gel is polymerized between the protein patterned glass coverslip and a silanized coverslip. (vi) After gel polymerization, the top coverslip is removed from the PAAm gel. Inset at right shows a fluorescence micrograph of a labeled protein transferred to a PAAm gel. (vii) Inset at right shows pairs of epithelial cells on the patterned PAAm gel restricting the geometry of the protein functionalized regions.

### Microcontact printing of ECM proteins (μCP)

We prepared PDMS stamps by casting Sylgard 184 PDMS (10:1 base to curing agent, Dow Corning) in a 9 μm deep mold microfabricated by standard photolithography using SU-8 negative resist ^11^. We incubated the PDMS stamps (1 cm^2^ squared stamps with 45 μm^2^ patterns with 80 μm spacing) with 100 μg/ml fluorescently labeled gelatin solution for one hour in the dark. Following protein incubation, we aspirated excess protein solution and dried the stamps gently using low nitrogen gas flow. Prior to μCP, we cleaned glass coverslips with 2% v/v Hellmanex solution (Hellma Analytics) in DI water for at least 30 minutes. We then rinsed the coverslips 5 times with DI water and dried them with compressed air prior to stamping. We put the PDMS stamps in contact with the cleaned coverslips for 5 minutes and removed the stamps by carefully forcing a tweezer between the coverslip and the edge of the stamp.

### Preparation of ECM patterned polyacrylamide gels

We transferred the protein patterns from the glass coverslip to the surface of PAAm gel for both the LOP and μCP protocols by co-polymerization (Figure 1, v-vii). Polyacrylamide gels of varying stiffness were polymerized between the protein patterned glass coverslip and a silanized bottom coverslip. The bottom coverslip was silanized to ensure covalent bonding of gels to this bottom glass layer, following a method by Guo and colleagues ^24^. Briefly, 30 μL of working solution (3 μl bind-silane (3-methacryloxypropyl-trimethoxysilane, Sigma-Aldrich, M6514), 950 μL 95% ethanol, and 50 μL of glacial acetic acid) were applied to the coverslip, allowed to incubate for 5 min, and then rinsed with ethanol and dried in a desiccator.

Polyacrylamide gels of three different stiffness were used for experiments: 5 kPa, 10 kPa, and 25 kPa as determined by Tse and Engler ^2^. MilliQ water, acrylamide (0.5 g/mL stock, Sigma-Aldrich, 01696 FLUKA), and bis-acrylamide (0.025 g/mL stock, Sigma-Aldrich, 146072) were combined to yield 5% w/v acrylamide and 0.15% w/v bis-acrylamide for 5 kPa gels, 10% w/v acrylamide and 0.1% w/v bis-acrylamide for 10 kPa gels, and 10% w/v acrylamide and 0.25% w/v bis-acrylamide for 25 kPa gels. The precursor solution was degassed in a vacuum desiccator for 1 hr. To initiate gelation, 5 μL of 10% w/v ammonium persulfate (APS, Sigma-Aldrich, A9164) was added to ~995 μL of gel precursor solution followed by 0.5 μL of N,N,N′,N′-Tetramethylethylenediamine accelerator (TEMED, Sigma-Aldrich, 411019). We missed the solutions by gentle pipetting, dispersed 50 μL of the solution on the activated coverslip, and then placed the protein-functionalized coverslip on top, creating a sandwich (Figure 1 v). Gels were left undisturbed at room temperature for 30 minutes to polymerize. After polymerization, the gels were immersed in PBS for at least 1 hour and the glass coverslip was removed from the top of the gels (Figure 1 vi).

### Analysis of Pattern Transfer Efficiency

To assess the protein transfer efficiency from the patterned glass coverslip onto the PAAm gel, we imaged the coverslips before transfer and the PAAm gel surface after gelation and coverslip removal using the same microscope image acquisition parameters. All acquired images were processed by ImageJ (http://rsb.info.nih.gov/ij/). We analyzed 150 individual patterned features by measuring the difference between the same feature on the coverslip before and after transfer, using the cvMatch_Template ImageJ plugin ^25^. The average background signal was determined outside the protein pattern and subtracted for each image. We measured the average pixel intensity within a region of interest defined as our theoretical patterning shape and calculated the transfer efficiency as the average intensity of the protein pattern on the gel image divided by the average intensity of the pattern on the coverslip.

### Analysis of the Geometric Accuracy of Protein Patterning

To compare the accuracy of patterns generated by LOP and μCP, we calculated the cross-correlation coefficient between the theoretical pattern shape and the binarized patterned features using the *corr2* function in Matlab (R2014b, Mathworks). The binarized stacks (n =150) were created with ImageJ by de-noising the images using the built-in despeckle function followed by automated binarization of each pattern using Otsu thresholding. Profile column average plots were analyzed from the binarized pattern stacks using ImageJ. To perform yield analysis, we selected around 389 – 416 features for each gel stiffness type, created a montage, and then used cross correlation with a threshold of 0.84 to find acceptable features. We divided the number of acceptable features by the total number of features analyzed for each gel stiffness to calculate the yield.

### Cell culture

Madin-Darby Canine Kidney (MDCK) type II G cells were transfected with LifeAct-GFP (ibidi, 60101) using the Amaxa Biosystem Nucleofector II system and transfection kit (Lonza, VCA-1005). The LifeAct-GFP MDCK cells were maintained in low glucose DMEM (Invitrogen, 11885) containing 1 g/l sodium bicarbonate, 1% Penicillin-Streptomycin (PenStrep, ThermoFisher, 15140122), 0.5 mg/ml G418 selection reagent (Sigma-Aldrich, G418-RO Roche), and supplemented with 10% (vol/vol) fetal bovine serum (FBS). 25 kPa PAAm gels patterned with 100 μg/ml collagen I (Gibco, A1048301) mixed with 20 μg/ml Alexa Fluor 568 labeled gelatin were cast into Mattek dishes (14 mm glass, Mattek P35G-0.170-14-C). MDCK cells were trypsinized and seeded on the PAAm gels for 16 hours before imaging experiments. Prior to imaging, the media was replaced to low glucose DMEM with no phenol red (ThermoFisher, 11054001) and supplemented with 1% PenStrep, 10% FBS, and 25 mM HEPES buffer. Cells were imaged on a Leica DMI3000B microscope with heated incubation unit at 5 minute intervals using a 40x air objective.

## Results

We compare the LOP and μCP methods by analyzing the efficiency of protein transfer from the surface of coverslips onto the surface of PAAm gels (Figure 2) and the geometrical accuracy of the created patterns (Figure 3). We use a square ‘frame’ pattern shape to compare how both protocols resolve corners and edges of a complex shape. We show pattern arrays of glass and PAAm samples normalized for contrast to aid visual comparison of the transfer efficiency for LOP and μCP techniques in Figure 2A and 2B.

**Figure 2.**
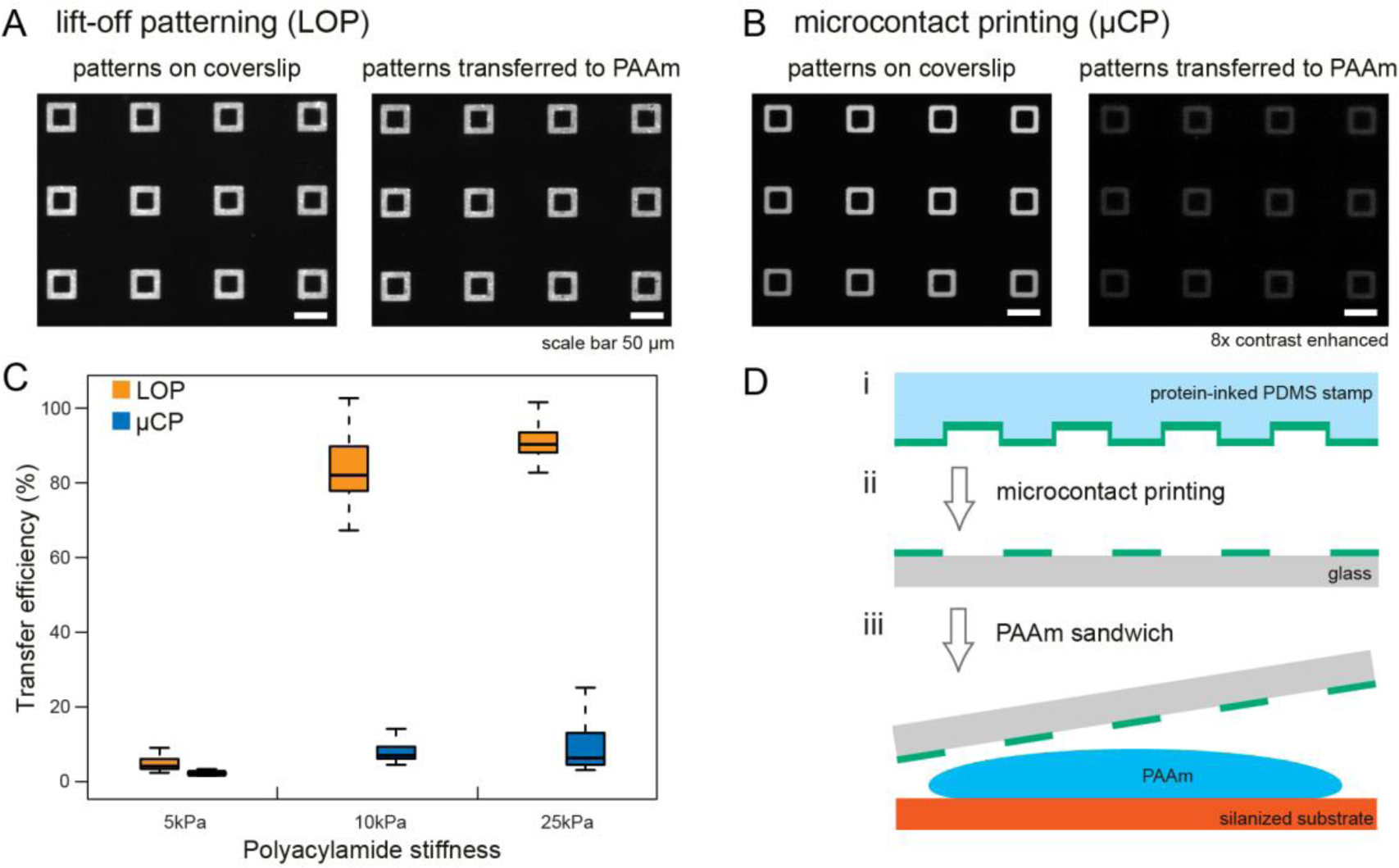
Quantification of protein transfer efficiency to PAAm gels of varying stiffness. (A,B) Arrays of 45 μm^2^ square protein patterns on 25 kPa PAAm gels created by LOP and μCP before and after transfer to gel surface. (C) Quantification of protein transfer efficiency from glass coverslips to PAAm gel of varying stiffness. Differences between LOP and μCP for each stiffness are statistically significant (p-value < 2.2E-16, Mann-Whitney-Wilcoxon test). Substantially more protein is transferred from patterns created by photoresist lift-off. Data are represented as box plots. The median, 1^st^ and 3^rd^ quartile, and minimum and maximum values are shown, n = 150 for each method and stiffness shown. (D) Overview of μCP method to pattern proteins on PAAm gels.

**Figure 3.**
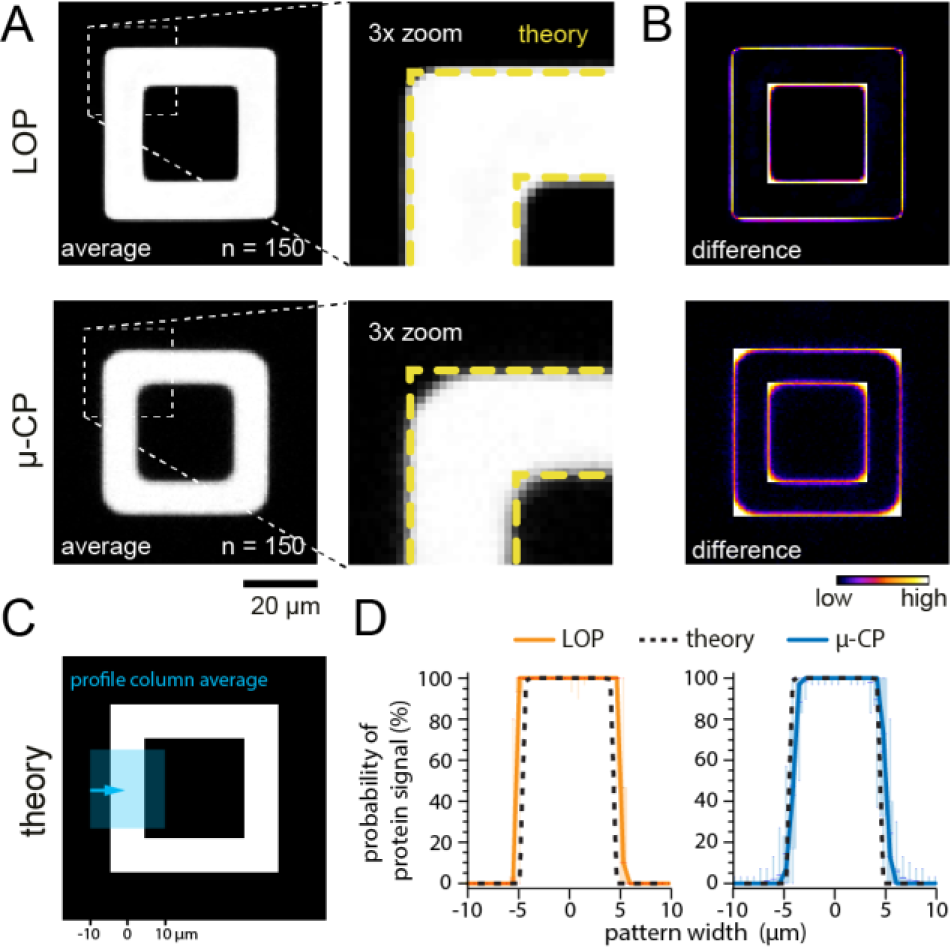
Comparison of pattern accuracy between LOP and μCP methods. (A) Average images of 150 binarized protein patterns created by LOP and μCP on 25 kPa gels. (B) Difference images calculated by comparing the average images and the theoretical pattern mask. Edges and corners are resolved substantially better in patterns created by LOP. (C) Theoretical pattern shape with a region highlighted corresponding to where profile column average scans were taken. (D) Profile column average scans across 150 binarized patterns show that the variation in protein signal at the pattern edges is strongly reduced in LOP patterns. Plotted are the median (line), 1^st^ / 3^rd^ quartile (box) and 5-95% (whisker) of the probability of protein present across the pattern width.

Protein patterns created by the LOP method are transferred more efficiently from the coverslips to gels for all gel stiffness we tested (Figure 2C). We find significant differences in transfer efficiency between LOP and μCP when comparing both protocols at each stiffness (p-value < 2.2E-16 using the Mann-Whitney-Wilcoxon to compare the 5 kPa, 10 kPa, and 25 kPa PAAm gel samples). However, the protein transfer efficiency in both methods is considerably lower for 5 kPa when compared to 10 kPa and 25 kPa PAAm samples. To explain this observation, we analyzed and compared the size properties of gelatin and polyacrylamide gel formulations used in this study to those commonly used in mechanobiology and electrophoresis (see Supplemental Information). The 10 kPa and 25 kPa gel formulations we use contain 10% total polymer which is twice that of the 5 kPa gel formulation (5%). Due to the lower total polymer content, we hypothesize that fewer sites are available for protein integration during polymerization in the 5 kPa gels. This effect is independent of the patterning technique and thus can be a limiting factor for the protein functionalization of soft polyacrylamide gels.

To evaluate the accuracy of the features transferred to PAAm gels, we compare the corners of the square frame patterns for both protocols (Figure 3A) and show the difference between the actual and theoretical shape (Figure 3B). Protein patterns created by LOP exhibit greater definition in the pattern edges and corners than protein patterns created by μCP. Cross-correlation analysis of the patterns compared to the theoretical pattern shape on the photomask shows that LOP results in patterns that more accurately recapitulate the theoretical shape. Correlation coefficients are as follows (n = 150 patterns; mean +/- standard deviation): μCP (5, 10, 25 kPa): 0.84±0.05; 0.87±0.02; 0.89±0.02; LOP (5, 10, 25 kPa): 0.91±0.04; 0.94±0.02; 0.93±0.01). The higher fidelity of the pattern edges becomes evident when we compare profile scans across the average of 150 patterns for both methods to the theoretical pattern shape (Figure 3C, D; similar to methods by Vignaud and colleagues ^9^). The variation in the protein signal at the pattern edges is strongly reduced in the LOP patterns. These results are also supported by a cross-correlation analysis where we tested the variability and yield of acceptable features across about 400 total feature samples for each gel formulation. We applied a correlation coefficient threshold for acceptable features of 0.84 to match the lowest correlation coefficient in our analysis above. LOP resulted in a greater number of acceptable features than μCP and acceptable feature yield varied from 59% to 98% for LOP and from 4% to 72% for μCP for different gel formulations (see SI Figure 2 and SI Table 1 for summary of data).

We demonstrate that single cells as well as pairs of cells attach exclusively to the ECM patterned areas of PAAm gels patterned using the LOP method (Figure 4A, B; SI Movies 1, 2). Areas between patterns exhibit anti-adhesive properties and prevent cells from binding outside the protein features. To test if the cytoskeletal architecture and remodeling are different for cells attached to patterns created by LOP or μCP, we followed the actin dynamics of LifeAct-GFP transfected MDCK cells using live cell fluorescence microscopy on 25 kPa substrates. We chose to conduct our analysis on gels with a 25 kPa elastic modulus because this value is close to the measured stiffness of a MDCK monolayer by micro-indentation: 33 ± 3 kPa ^26^. Consistent with previous literature reports ^27^, we found that cell doublets on the frame patterns rotated around each other (Figure 4B,D; SI Movies 2, 4). While this was observed independent of the patterning method used, the cell edges were more clearly defined for cells adhering to patterns created by LOP than gels patterned by μCP (Figure 5), which was consistent with the higher pattern accuracy (Figure 3).

**Figure 4.**
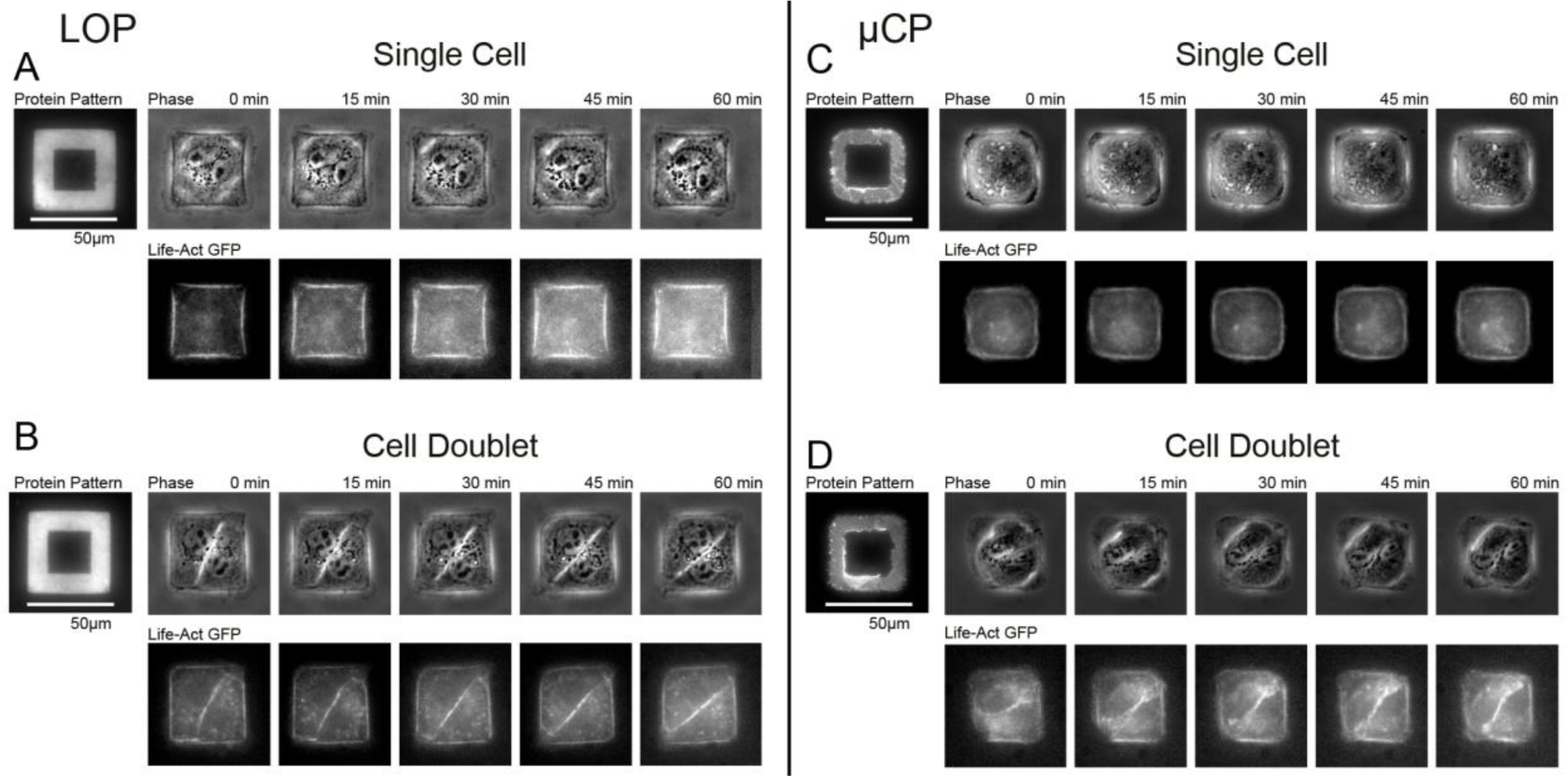
LOP yields sharper cell edges with localized actin bundles compared to μCP patterned gels. Time-lapse acquisitions of MDCK cells transfected with Lifeact-GFP (actin label) grown on 25 kPa PAAm gels showed similar intracellular actin structures on LOP (A,B) and μCP (C,D) protein patterns. Cell doublets rotated around each other on the patterns for both techniques (B,D).

**Figure 5.**
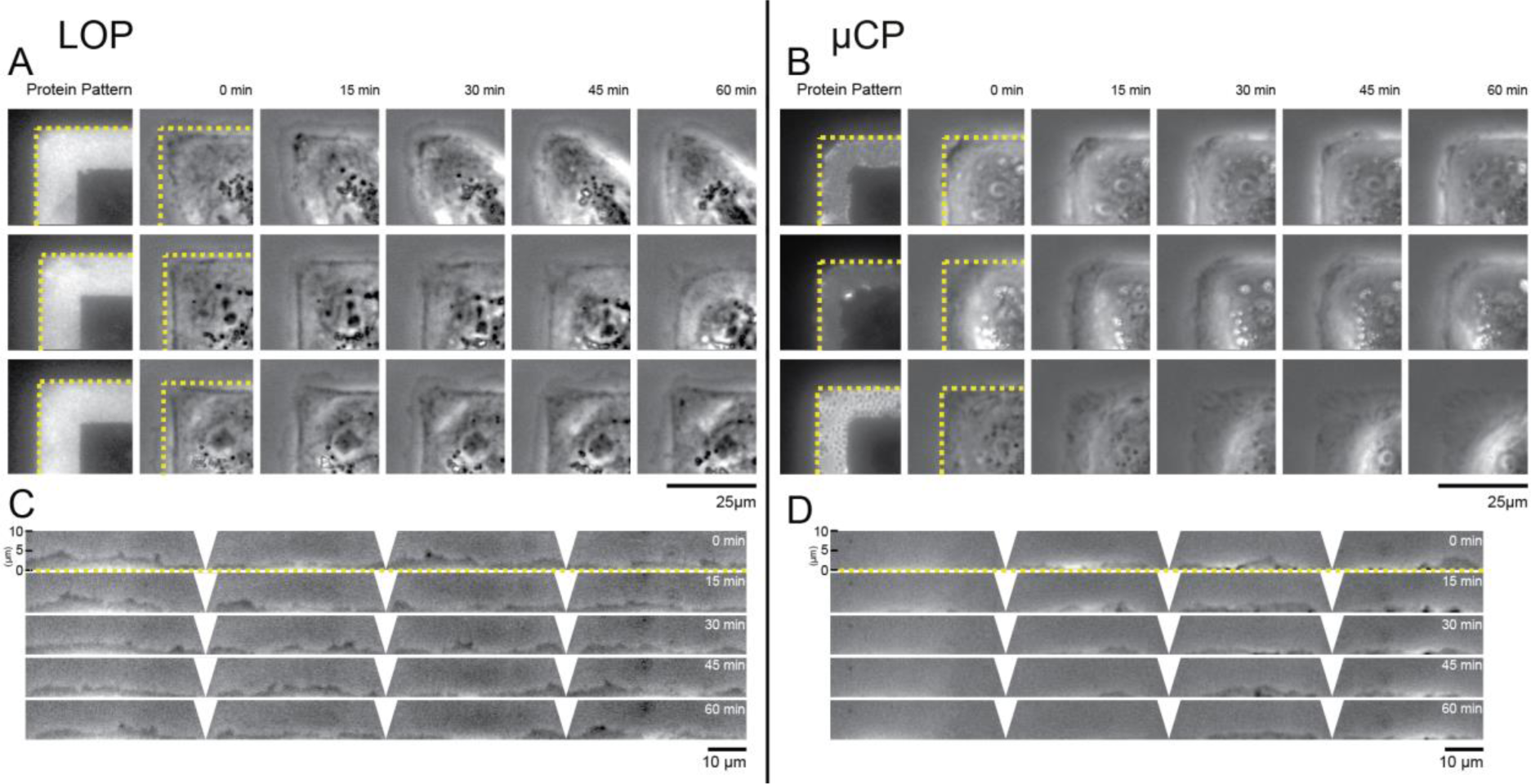
Lamellipodia are more exploratory for cells on LOP than on μCP patterned substrates. Phase contrast time lapse imaging of 3 representative MDCK cells on 25 kPa PAAm gels patterned by LOP (A) and μCP (B). Cells on substrates produced by LOP follow the protein pattern border more accurately (dotted line panels A, B) and reveal more pronounced lamellipodia. (C, D) Kymograph analysis of lamellipodia kinetics along pattern edge. Single cell on LOP pattern shows increased lamellipodia protrusions and retractions within a 10 μm wide ROI outside the protein pattern edge compared to a cell on a μCP pattern (SI Movie 5,6). Kymographs show cells depicted in bottom row of panel A and B. ROIs are straightened and distorted regions at the pattern corners are cleared.

## Discussion

In this work, we introduce a photoresist-based LOP technique to pattern ECM proteins on polyacrylamide hydrogels to control the shape of cells with high-fidelity and compare it with the widely used μCP protocol. We found the LOP method to be more efficient and accurate in reproducing complex micrometer-sized patterns (Figure 2,3). To illustrate that the improved fidelity of LOP patterns translates to greater control over cell shape, we cultured MDCK epithelial cells on patterned 25 kPa gels for up to 16 hours. The shape of single cells and cell pairs on LOP pattern reflected the theoretical shape with greater accuracy as compared to cells on μCP patterns (Figure 4,5; SI Movies 1 - 4).

The differences in pattern fidelity on the hydrogels stem from differences in the patterning of the glass coverslips. LOP relies on the molecular assembly of the biopassive PLL-g-PEG copolymer on S1818 photoresist features directly on the glass substrates while μCP involves PDMS replica molding from SU-8 master structures and stamping of the protein to the coverslip. The ideal spatial resolution that is achievable by contact photolithography can be estimated using the following relation between the exposure wavelength (*λ*) and the photoresist thickness (*z*) ^28^:

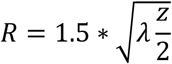

The S1818 resist for LOP is 2μm thin compared to the 9μm SU-8 photoresist layer used as master structure for fabricating PDMS stamps. Thus, the difference in resist thickness substantially contributes to a decrease in pattern accuracy for μCP. Deep reactive ion etching (DRIE) of silicon substrates patterned with a thin photoresist layer can be used to circumvent photolithography of thick resist layers for the fabrication of the master structures for PDMS replica molding^29^. The height of the master structures can be controlled by the etching time, yet this process requires specialized equipment not readily available in many laboratories.

In addition to the resist layer thickness, the type of resist used in both protocols contributes to pattern fidelity. The negative SU8 resist used in μCP to create the PDMS mold is typically underdeveloped and thus resolving the edges of features is a challenge (see Supplementary Information, SI Figure 1) which explains why μCP results in features which are smaller than the theoretical specifications (see Figure 3A). LOP uses the positive S1818 resist which tends to be over-developed and thus features fabricated by this method tend to be larger than theoretical specifications but with better-resolved corners (see Figure 3A). Further, LOP enables the design and fabrication of arbitrary pattern geometry and spatial organization (e.g., large pattern-to-pattern distance) as LOP circumvents the need to maintain specific height-width ratios of the PDMS stamps to keep the elastomeric structures from collapsing during μCP ^14, 30^. Additional sources for error in μCP can arise from the drying of proteins on the PDMS step prior to transfer and the non-uniform contact of the stamp with the glass coverslip.

We found the lamellipodia of MDCK cells to be more exploratory and dynamic on LOP than μCP substrates, extending up to 5 μm outside of the protein pattern (Figure 5). Epithelial cells have been shown to extend lamellipodia for several micrometers past areas with ECM during wound healing and cell migration ^31, 32^. Our observations of lamellipodia extending up to 5 μm beyond adhesive regions and the pronounced actin bundles at the pattern edges match well with the spatiotemporal organization of the cytoskeleton and focal adhesions at the leading edge of migrating cells ^33^. We were interested to note this difference in cell exploration and we expect the LOP method to enable future studies on the role of ECM organization on cell migration and lamellipodia extension.

An open question for patterns created by LOP is whether the PLL-g-PEG blocking agent on the glass coverslips is transferred to the PAAm gel. We are not able to trace fluorescently labeled PLL-g-PEG molecules in the gel samples but individual PEG chains have been shown to polymerize into the backbone of polyacrylamide when added in bulk to the precursor solution ^34^. The interactions of PLL-g-PEG copolymer and polyacrylamide during gel polymerization are unclear. Regardless of PLL-g-PEG transfer to the PAAm surface, LOP results in functionalized PAAm gels with nonadhesive regions between protein patterns. We noted that removing the glass coverslips from polymerized gels was easier for samples created by LOP than for μCP (step vi, Figure 1) and we hypothesize that this effect could be due to the high water content of the PLL-g-PEG layer on the coverslips ^35^.

**In summary,** our LOP method facilitates advanced cell culture techniques that require precise patterning of single or multiple cells into a variety of shapes on hydrogel substrates of varied stiffness. High pattern accuracy and defined ECM density within the protein patterns are essential to compare cell phenotypes on different patterns and reduce the systematic error of pooled measurements. This is of particular importance for studies focusing on complex, multivariate cell-ECM signaling pathways and the cytoskeletal response to different cell geometries and substrate stiffness ^36^. Overall, local ECM density, cell shape, and substrate stiffness have been shown to regulate the structural organization of focal adhesion complexes ^37, 38^, the force balance between cell-cell and cell-ECM adhesions ^4^, the nuclear lamina ^39^, mesenchymal stem cell stiffness ^40^, stem cell fate ^12, 41^, and the contractile properties of cardiomyocytes ^42^.

## Acknowledgements

The authors thank Pruitt lab members for helpful discussion of results. This project was supported in part by the National Science Foundation (EFRI-MIKS 1136790,), National Institutes of Health (R01EB006745, 1R21HL13099301). The authors are grateful for graduate and post-doctoral research fellowships from the National Science Foundation, ILJU Foundation, Stanford BioX, Stanford Office of the Vice Provost for Graduate Education, American Heart Association, and Stanford Graduate Fellowship.

